# Unsupervised cluster analysis of SARS-CoV-2 genomes indicates that recent (June 2020) cases in Beijing are from a genetic subgroup that consists of mostly European and South(east) Asian samples, of which the latter are the most recent

**DOI:** 10.1101/2020.06.22.165936

**Authors:** Georg Hahn, Michael H. Cho, Scott T. Weiss, Edwin K. Silverman, Christoph Lange

## Abstract

Research efforts of the ongoing SARS-CoV-2 pandemic have focused on viral genome sequence analysis to understand how the virus spread across the globe. Here, we assess three recently identified SARS-CoV-2 genomes in Beijing from June 2020 and attempt to determine the origin of these genomes, made available in the GISAID database. The database contains fully or partially sequenced SARS-CoV-2 samples from laboratories around the world. Including the three new samples and excluding samples with missing annotations, we analyzed 7, 643 SARS-CoV-2 genomes. Using principal component analysis computed on a similarity matrix that compares all pairs of the SARS-CoV-2 nucleotide sequences at all loci simultaneously, using the Jaccard index, we find that the newly discovered virus genomes from Beijing are in a genetic cluster that consists mostly of cases from Europe and South(east) Asia. The sequences of the new cases are most related to virus genomes from a small number of cases from China (March 2020), cases from Europe (February to early May 2020), and cases from South(east) Asia (May to June 2020). These findings could suggest that the original cases of this genetic cluster originated from China in March 2020 and were re-introduced to China by transmissions from samples from South(east) Asia between April and June 2020.

## 1 Introduction

One important aspect being investigated about the ongoing pandemic of the SARS-CoV-2 virus pertains to its geographic progression around the world and the linked question of possible viral mutations during viral spread.

We focus on the recent report of three newly diagnosed cases in Beijing and their viral genomes that were reported in June 2020. One working hypothesis is that these new cases originated from Europe (Xiaohua, 2020; Liu and Nebehay, 2020). We utilized an unsupervised clustering approach where results of the previously sequenced viral genomes matched the history of the geographic spread of SARS-CoV-2 (Hahn et al., 2020a).

The approach we employ is an unsupervised clustering technique considered in Hahn et al. (2020a). In this paper, we consider the computation of a similarity measure on the Hamming matrix of the SARS-CoV-2 genome samples available in the GISAID database (Elbe and Buckland-Merrett, 2017; Shu and McCauley, 2017). The Hamming matrix encodes the difference of all samples compared to the reference genome, and the similarity measure being employed is the Jaccard matrix (Prokopenko et al., 2016; Schlauch et al., 2017). Visualizing the first two principal components of the Jaccard similarity matrix revealed that the samples from North America and Europe fall into four branches. Those branches originated from the origin, which contained samples from Wuhan, other parts of China, and geographically proximate areas such as Southeast Asia, thus reflecting the geographic progression of the virus across the globe.

In this paper we repeat the PCA (principal component) analysis of the Jaccard similarity measure on the Hamming matrix of Hahn et al. (2020a). The dataset under investigation is a recent timestamp (14 June 2020) of single stranded RNAs of SARS-CoV-2 viruses, extracted from SARS-CoV-2 patients around the world and downloaded from the GISAID database. We contrast the three newly found SARS-CoV-2 genomes in Beijing, available on GISAID under the accession numbers *EPI_ISL_469254, EPI_ISL_469255* and *EPI_ISL_469256*, to the other 7, 640 SARS-CoV-2 patients without missing entries available in the database. The first two samples, *EPI_ISL_469254* and *EPI_ISL_469255*, are extracted from human patients, whereas *EPI_ISL_469256* is an environmental sample. We show that all three samples indeed fall into one of the European and South(east) Asian branches of the SARS-CoV-2 genome reported in Hahn et al. (2020a), which also contain a small number of older cases from China stemming from March 2020. As the most recent cases in these branches are almost exclusively from South(east) Asia, this could suggest that the new cases in Beijing were re-introduced by transmissions from South(east) Asia.

## 2 Methods

We downloaded all complete nucleotide sequences (defined as having a length ≥ 29, 000) of the SARS-CoV-2 genome available from the GISAID database (Elbe and Buckland-Merrett, 2017; Shu and McCauley, 2017) with a timestamp up to 14 June 2020. This dataset contains 45, 895 different sequences.

We proceed by keeping only those samples having a valid collection timestamp, and filter for sequences without reading errors, that is without missing base pair entries. Afterwards, we group samples into 13 distinct categories using their geographic location tag. Those categories were: (1) China (all provinces apart from Wuhan), (2) Wuhan, (3) Korea, (4) Europe (all countries apart from Italy), (5) Italy, (6) Africa, (7) Australia and New Zealand, (8) South(east) Asia, (9) North America (USA, Canada), (10) Central America (including Mexico), (11) South America, (12) Japan, (13) Iran.

We employ the anchor sequences of (Hahn et al., 2020a, Section 2) to align all sequences. Samples without a match of the anchor sequence were discarded, leaving *n* = 7, 643 samples for analysis including the new samples from China.

After aligning with the anchor, all samples as well as the SARS-CoV-2 reference sequence (available on GISAID under the accession number *EPI_ISL_412026*) were aligned in order to establish a window in which all sequences had reads. This window comprised a length of *p* = 28, 784.

We are interested in measuring the similarity between the samples in order to group them. Instead of comparing the samples directly, to have a common ground for comparison, we compare their differences to the reference sequences.

Similarly to Hahn et al. (2020a), we denote with *X* ∈ 𝕊^*n*×*p*^ the matrix of all *n* nucleotide sequences, where 𝕊 ∈ {*A, C, G, T*}. We denote with *r* ∈ 𝕊^*p*^ the reference genome from GISAID. We compute the Hamming matrix *Y* ∈ {0, 1}^*n*×*p*^ encoding the differences of each nucleotide sample compared to the reference sequence after alignment. Precisely, we define *Y*_*ij*_ = 1 if and only if *X*_*ij*_ = *r*_*j*_ for sample *i*, and *Y*_*ij*_ = 0 otherwise. Each entry *Y*_*ij*_ = 1 thus indicates a mismatch at position *j* of the sample *i* with the reference sequences. The row sums of *Y* are the Hamming distance of each sample to the reference sequence.

The resulting Hamming matrix *Y* is analyzed with the *locStra* methodology of Hahn et al. (2020b), using the *locStra* R-package on CRAN (Hahn et al., 2020c). In particular, we compute the Jaccard similarity matrix of *Y* (Prokopenko et al., 2016; Schlauch et al., 2017), *denoted Jac*(*Y*) ∈ ℝ^*n*×*n*^. The entry *Jac*(*Y*)_*ij*_ is the set theoretic similarity measure between the rows *i* ∈ {1, *…, n*} and *j* ∈ {1, *…, n*} of the Hamming matrix *Y*. The Jaccard matrix *Jac*(*Y*) is thus the matrix of all pairwise similarity measures among the *n* samples in GISAID. Computing the first two PCAs of *Jac*(*Y*) allows us to visualize the clustering.

Regarding the newly found samples in Beijing, we observe that all three samples *EPI_ISL_469254, EPI_ISL_469255*, and *EPI_ISL_469256* remain after filtering and can be aligned to the reference sequence.

## 3 Results

Figure 1 plots the first two PCAs of the Jaccard similarity matrix *Jac*(*Y*). All samples are color coded by their geographic tag in the GISAID database. Importantly, the samples from China are emphasized with large crosses. The two genome samples from the newly found human cases in Beijing, *EPI_ISL_469254* and *EPI_ISL_469255*, are visualized with large triangles. The recent environmental sample *EPI_ISL_469256* is projected on top of *EPI_ISL_469254* (the one closer to the origin).

**Figure 1:**
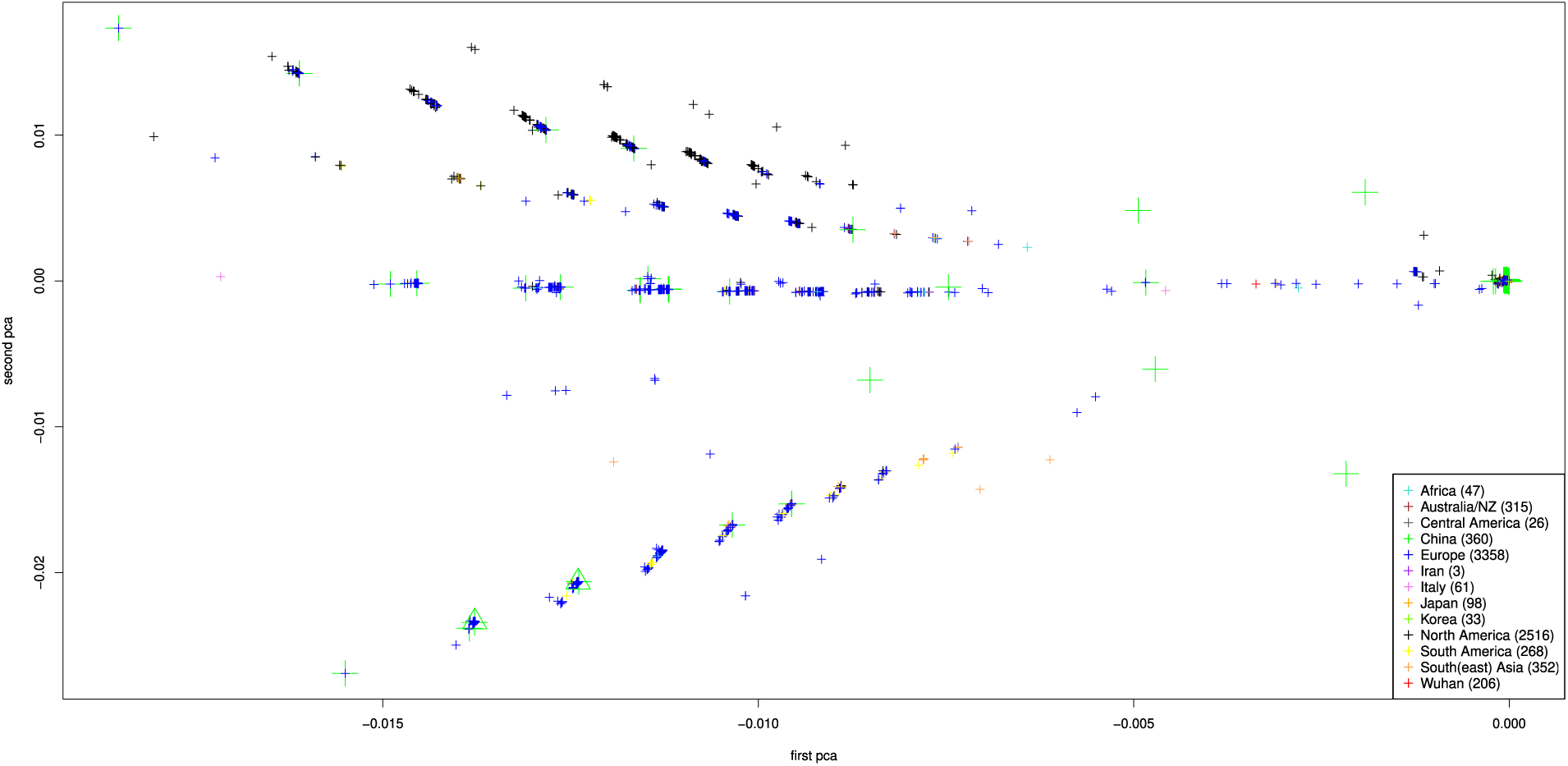
First two PCAs of the Jaccard matrix of *Y* (see Section 2. Colors indicate the geographic origin of the genome samples. The samples with geographic tag “China” are visualized with large crosses. The three newly found samples *EPI_ISL_469254, EPI_ISL_469255*, and *EPI_ISL_469256* are visualized with large triangles.

Three observations are worth noting. First, the overall clustering visible in Figure 1 is consistent with the one reported in Hahn et al. (2020a) despite having three times as many genome samples stemming from far more geographic regions (2, 540 samples are used in Hahn et al. (2020a), whereas here we use *n* = 7, 643; moreover, no African or Central American groups were considered in Hahn et al. (2020a)). The observations made in Hahn et al. (2020a) hold true: Starting from the origin containing most samples from China and nearby regions, we observe four branches containing North American, European and South(east) Asian samples. The top branch predominantly contains North American samples. European samples are mixed with North American ones in the second (from the top) branch, whereas both lower branches mostly contain European and South(east) Asian samples.

Second, samples originating from China are mixed throughout the four branches, especially the lower two branches containing mostly European samples, illustrating the spread of the disease out of Wuhan.

Third, the three new genomes under consideration here (*EPI_ISL_469254, EPI_ISL_469255*, and *EPI_ISL_469256*) fall into the lower branch of European and South(east) Asian samples, which also include a small number of cases from China in their vicinity (defined as having a Euclidean distance of at most 10^−4^ in the projected plane spanned by the first two principal components of the Jaccard similarity matrix). The cases from China in this cluster have timestamps going back to March 2020 (see Figure 2). While the majority of the European cases are from February to early May 2020, the cases from South(east) Asia belong to the most recent time period from May to June 2020. This could suggest that the new cases in Beijing were caused by transmissions that are linked to cases from South(east) Asia.

**Figure 2:**
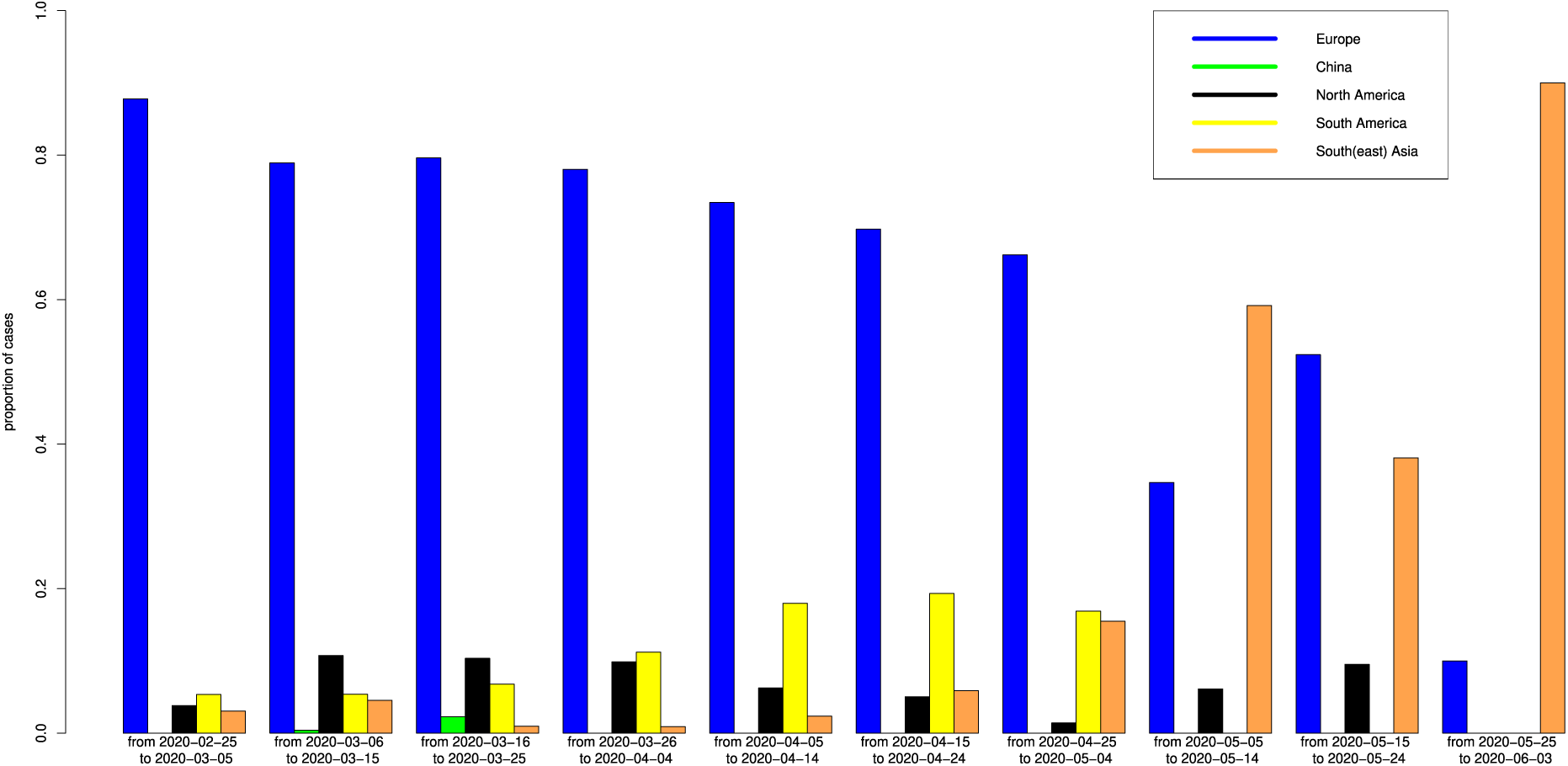
Proportion of cases (normalized per bin) in the vicinity of *EPI_ISL_469254* (defined with radius 10^−4^) as a function of time. The corresponding figure for the vicinity of *EPI_ISL_469255* is virtually identical and thus omitted here.

## 4 Discussion

We repeated an unsupervised clustering approach of SARS-CoV-2 samples available on the GISAID database, using the Jaccard similarity index computed on the matrix of Hamming differences between the nucleotide sequences of all genome samples and the reference genome. The original analysis performed in Hahn et al. (2020a) contained fewer samples, and those came from fewer geographic regions.

Here, we were particularly interested in three newly found samples with GISAID accession numbers *EPI_ISL_469254, EPI_ISL_469255*, and *EPI_ISL_469256* which were discovered in Beijing in June 2020. One working hypothesis is that the newly found cases were “imported” from outside. While the new cases do not form a new cluster, they fall into distinct clusters of cases that, with the exception of a few early cases from China (March 2020), are predominantly European and South(east) Asian cases. According to the sequencing time stamps, the most recent cases in these clusters are from South(east) Asia, which could imply that the new cases in Beijing are attributable to transmissions from South(east) Asian cases.

## Acknowledgements

The authors gratefully acknowledge the contributors, originating and submitting laboratories of the sequences from GISAID’s EpiCoV™ Database (Elbe and Buckland-Merrett, 2017; Shu and McCauley, 2017) on which this research is based. A detailed list of contributors is available in the Supplementary Information.

## Data Availability Statement

Sequence data that support the findings of this study are deposited in the GISAID database with accession numbers in the range of EPI_ISL_402119 to EPI_ISL_469256 (https://www.gisaid.org/).

## Funding

The initial methodology work for this paper was funded by Cure Alzheimer’s Fund; Funding for this research was provided through the National Heart, Lung, and Blood Institute [U01HL089856, U01HL089897, P01HL120839, P01HL132825] and National Human Genome Research Institute [R01HG008976].

## Notes

### Competing Interest Statement

The authors have declared no competing interest.

